# Sketch the Organoids from Birth to Death – Development of an Intelligent OrgaTracker System for Multi-Dimensional Organoid Analysis and Recreation

**DOI:** 10.1101/2022.12.11.519947

**Authors:** Xuan Du, Wenhao Cui, Jiaping Song, Yanping Cheng, Yuxin Qi, Yue Zhang, Qiwei Li, Jing Zhang, Lifeng Sha, Jianjun Ge, Yanhui Li, Zaozao Chen, Zhongze Gu

## Abstract

Organoids are three-dimensional *in vitro* models that recreate the structure and physiology of their source organs or tissues in remarkable detail. Due to the diversity of organoids in shape and size and the three-dimensional growth environment, it is challenging to observe and analyze organoids periodically in the microscope to obtain their morphological or growth characteristics, especially in high-throughput experiments. Here, this paper first proposes OrgaTracker, a novel assembled architecture combining Yolov5 for object detection and U-net for semantic segmentation. The deep learning algorithm can track and segment organoids over time and effectively avoid the influence of bubbles and accurately capture and analyze organoid fusion. A two-stage object detection methodology was performed to achieve the crypt count of each mouse small intestinal organoid, and the skeleton of intestinal organoids was further extracted to describe the structural relationship between the absorption villi and the crypt. Next, we used the “sketch” to convey visual concepts, which led to the clear identification of organoids at different growth/treatment stages. Lastly, based on our designed GAN network, various organoid images could be generated by drawing sketches, which for the first time provided a direct and practical approach for biologists and researchers to create “artificial organoids” simulating organoid morphology and allowing the exchange of ideas on organoid development. In sum, this research reported and provided a comprehensive novel organoid analysis and generation methodology for organoid research.

**Highlights:** OrgaTracker can track organoids and capture and analyze the integration of organoids. The system was also able to identify the number of crypts in each intestinal organoid, as well as extract the skeleton of the organoid. It also allowed, for the first time, recreating “artificial organoids” from hand-drawn sketches.

## Introduction

Currently, more than 80% of clinically tested drugs fail to pass clinical trials, with the vast majority of these failures being due to toxicity and lack of efficacy. [1]. To address these issues and provide alternative approaches for studies at the preclinical stage, human-cell/tissue-derived organoids are becoming increasingly favored in vitro research tools by biologists, medical researchers, and engineers [2]. Organoids mirror in vivo tissue organization and are powerful tools for investigating cell development and biology. Researchers have built models that simulate intestinal infections using microinjections of mouse intestinal organoids, allowing direct observation and examination of pathogen interactions with primary epithelial cells [3]. In addition, organoids have been used to create new model systems for studying human development and disease, e.g., cultured brain organoids have been used to demonstrate neural differentiation, cortical organization, and zonation, reproducing critical features of the early stages of human brain development [4, 5].

Different organoids have specific structures according to their original organs [6-9]. The morphology and structure of organoids can also reflect their growth/developmental state, e.g., the area of colorectal cancer organoids can show their viability; the number of buds in intestinal organoids represents their maturity; the textures of cerebral organoids, such as rosette structures, reflect the neurons’ differentiation and maturity; etc. However, there are very few automated tools for the analysis of organoids, and the judgment of whether organoids are growing well or not still mostly depends on scientists’ visual analysis and the results of complex immunofluorescence staining processes that are applied after sectioning. There is an urgent unmet need for effective analytical approaches, especially for the quantitative analysis and evaluation of organoids, to reduce the subjective error and heavy workload associated with human visual observation in focusing and searching operations.

Three-dimensional imaging and analysis of organoids faces the following obstacles. Unlike 2D cell cultures, organoids are usually expanded and grown in 3D environments using naturally derived or synthetic extracellular matrices. Assessing these cultures’ morphology and growth characteristics has been difficult for the following reasons: (I) Various imaging artifacts. Occluding or overlapping organoids and partially or totally out-of-focus of organoids pose many challenges for AI recognition [10, 11]. (II) Exclusion of air bubbles and unwanted dust. Bubbles and dust are likely to occur when Matrigel is added to the well plate to form a dome or when the culture medium is changed. A pipette touching the bottom of the well plate, plate shaking, and incorrect operations also frequently lead to bubbles [12]. (III) The complexity of 3D compositions containing multiple organoids. Some scientists have used cerebral organoids to assemble multiple different brain regions to create “mini brains” [13]. (IV) Organoid destruction and fusion. Other scientists have modeled human hepato-biliary-pancreatic organogenesis using a multi-organoid destruction and fusion approach [14]. For these reasons, many existing cell analysis tools, such as algorithms for high-content cell imaging and analysis and algorithms for immunohistochemistry images, cannot be applied to the analysis of organoids [15, 16]. We recently reviewed research papers on organoids combined with AI algorithms and found that there were not many related articles. The existing articles realized only simple recognition and segmentation of organoids, and the algorithms used lagged far behind the development of AI [17, 18]. Our team has been working on constructing organoids and organ-on-a-chip systems for over ten years, as well as tools for the intelligent identification and analysis of their images [19-23]. In recent studies, we used U-Net to segment tumor spheres to further quantify their invasiveness [24] and VGG to identify the responses of epithelial and endothelial cell layers and macrophages in a lung-on-a-chip to different stimuli [25]. The intelligent recognition of organoids needs more detailed analysis and further research.

In this study, we constructed tumor organoids, mouse small-intestinal organoids, and human cerebral organoids, and developed intelligent algorithms to analyze these 3D-cultured organoids (Fig. 1a-b). Our deep-learning algorithms were able to perfectly remove bubbles and impurities in the images; recognize the growth, fusion, and death of organoids after drug treatment; and carry out analysis of organoid characteristics, including budding, morphology, and skeleton (Fig. 1c-d). Finally, we realized for the first time intelligent generation of organoid images mimicking tumor organoids, small-intestinal organoids, and cerebral organoids (Fig. 1e). We believe that this work could provide a comprehensive scientific methodology and tool for researchers in biology, medicine, and engineering who are interested in vitro organoids and organs-on-a-chip.

**Figure 1.**
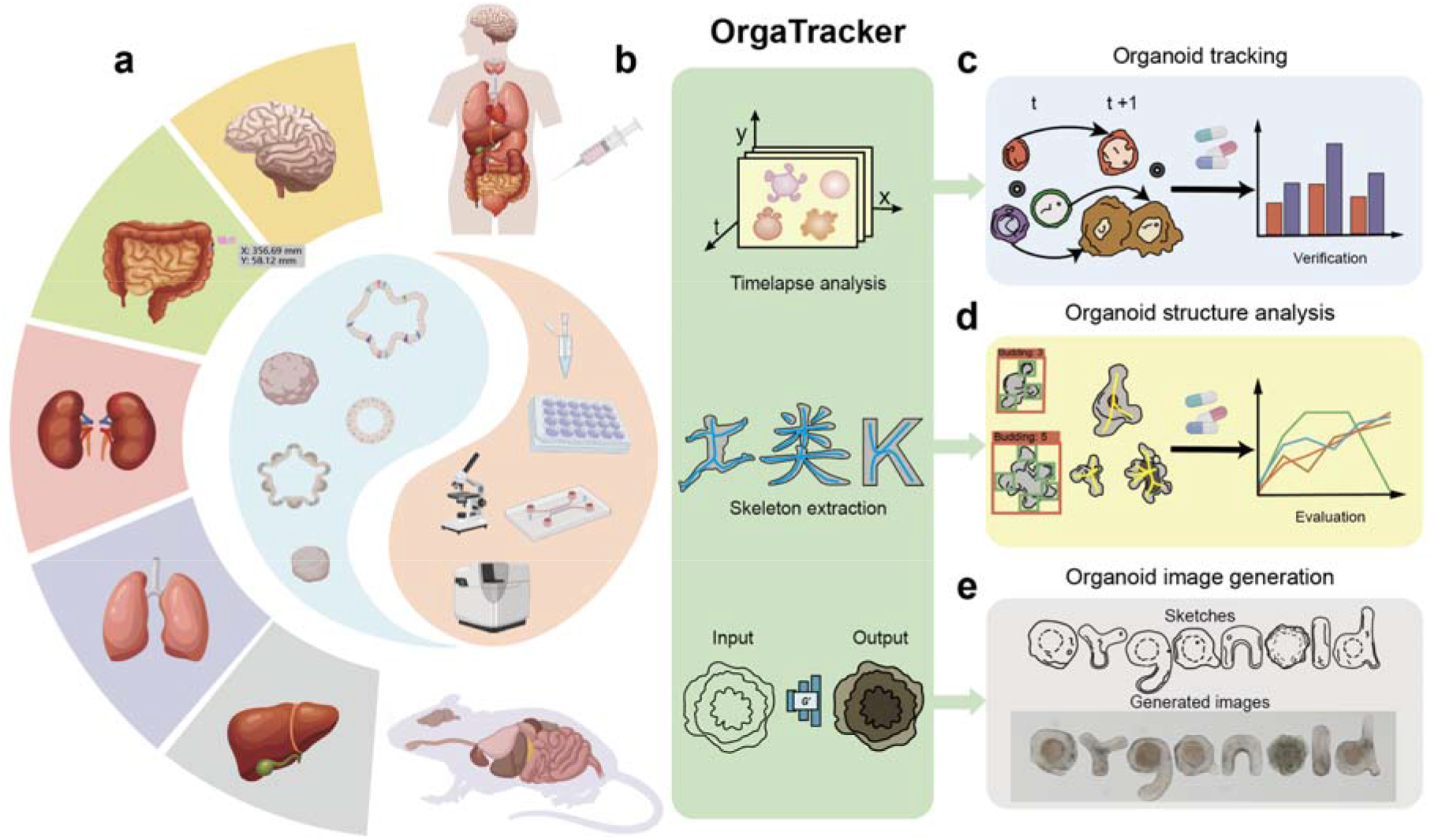
The Overall Flow of an Intelligent OrgaTracker System for Multidimensional Organoid Analysis. **a**. Organoid collection, culture, observation, imaging. Tumors, intestinal and cerebral organoids were collected from human or mouse bodies and cultured, observed and imaged under certain conditions. **b**. The input and composition of OrgaTracker. The input data are the images of various organoids under the bright-field, and the network included convolutional neural network, digital image processing and generation adversarial network, combing Image processing can help us extract the skeleton of all kinds of things, like football players, Chinese characters, the letter k. **c**. Organoid tracking and analysis. The system can realize tumor organoid matching at adjacent time intervals, filter bubbles, capture tumor organoid fusion and analysis after drug. **d**. Organoid structure analysis. The algorithm can realize budding count, skeleton extraction, intestinal organoids and analysis after drug. **e**. Organoid image generation. Based on GAN, the algorithm used sketches to generate the word “organoid” from the mouse small intestinal organoids and cerebral organoid.

## Results

### I. Overall Principle and Model Establishment

An overview is shown in Figure 2a of our proposed OrgaTracker architecture, which can achieve organoid tracking and area analysis under high-throughput imaging conditions. First, the improved Yolov5 is employed for organoid detection and location. Second, the detected regions and location information are sent into the improved U-Net to achieve the segmentation of organoids. The OrgaTracker is initialized using the first results of the object detection and assigns each organoid an ID starting from 0. Third, *IOU* matching is performed on organoids in the adjacent frame images output by these networks. Then, the Hungarian algorithm uses the IOU-matching values of different objects in adjacent frames to achieve optimal matching. The matched objects are assigned the previous ID, and the mismatched objects are assigned a new ID. Finally, the result is postprocessed and output to the image. We introduce the Transformer framework in the field of natural language processing to form the CNN and Transformer architecture as the Yolov5 backbone network, and make full use of the advantages of CNN and Transformer to improve the object detection accuracy (Fig. 2b). In addition, replacing NMS with DIoU-NMS can improve the detection accuracy of overlapping and occluded objects. Referring to the structure of U-Net and U-Net++, some skip connections are added between the U-Net encoder and decoder to integrate multiscale image features of different layers, which can improve the segmentation accuracy of organoids (Fig. 2c). These models were trained using the tumor organoid dataset. After verification and testing, our modified Yolov5 and U-Net’s performance was improved (Fig. 3a-c).

**Figure 2.**
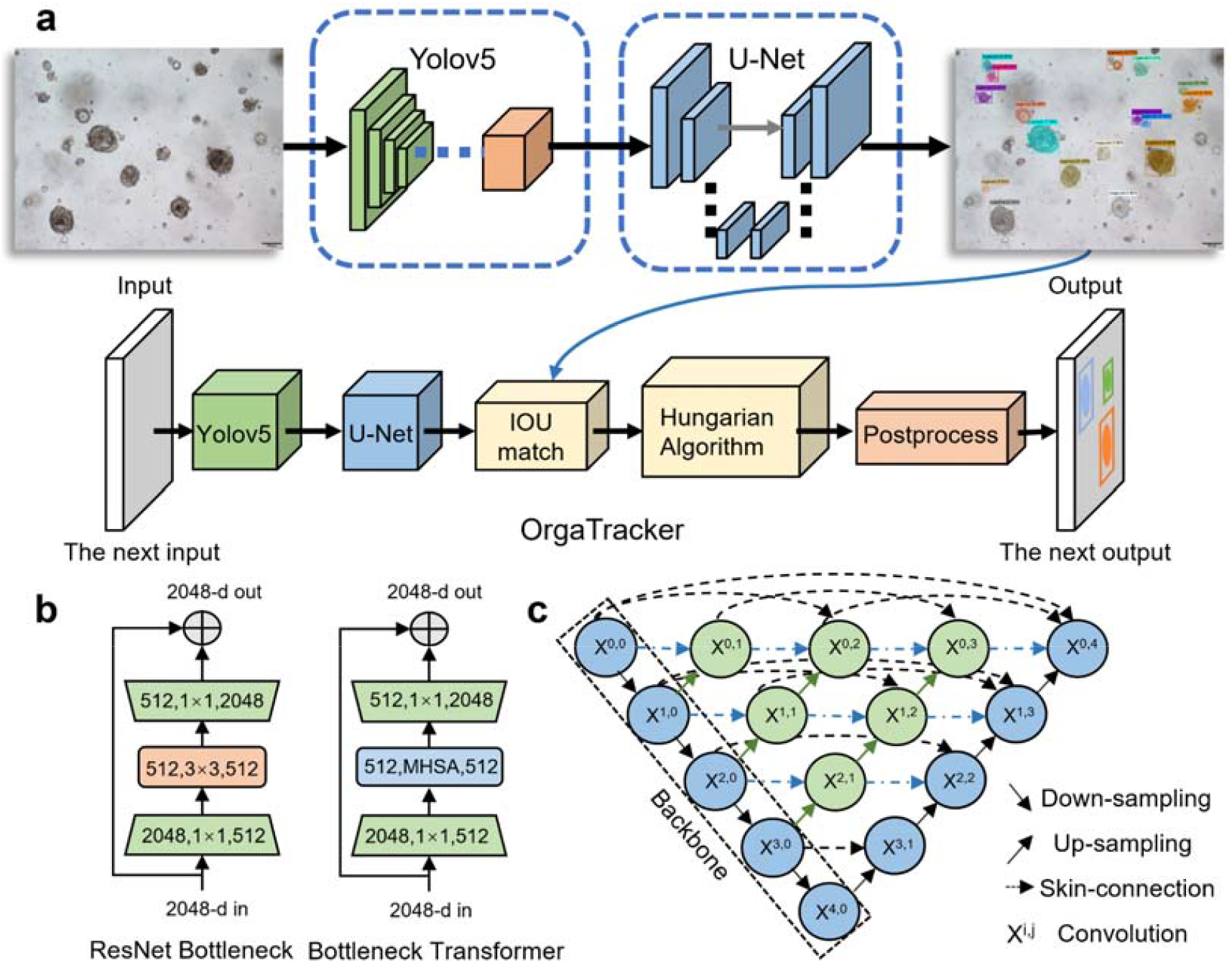
The Algorithm Flow of Organoid Tracking and Improved Network Architecture. **a**. The algorithm flow of OrgaTracker. The Yolov5 network is used to detect and locate organoids, and then the detected results are transmitted to U-Net for organoid segmentation. The organoids of adjacent images processed by the models are matched by IOU and the Hungarian algorithm to track the organoids. **b**. The improvement of Yolov5 backbone. The backbone network replaces ResNet Bottleneck with Bottleneck Transformer to form the architecture of combining CNN and Transformer. **c**. The improvement of U-Net backbone network. The backbone network integrates more features of different layers to improve the network segmentation ability.

**Figure 3.**
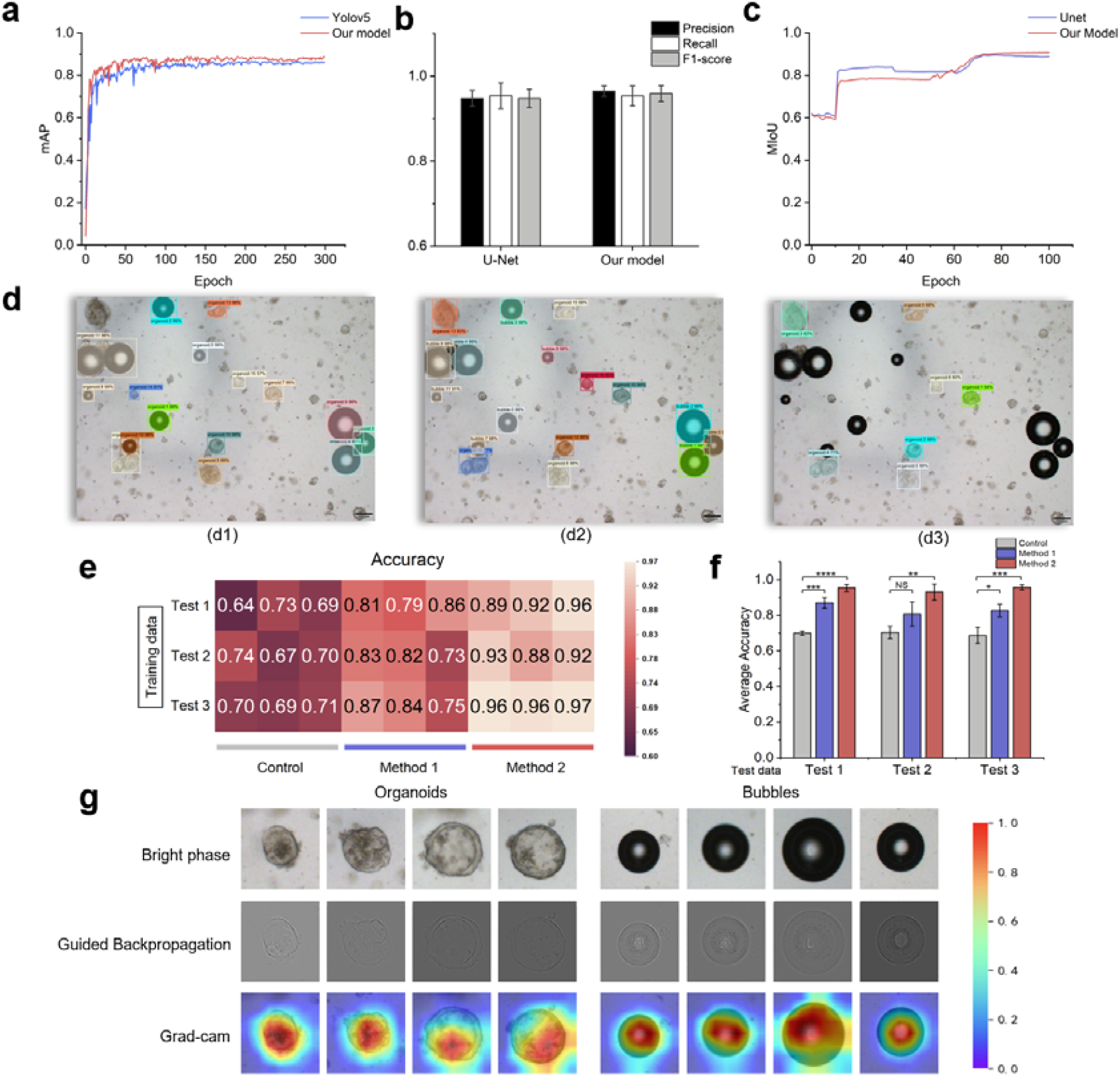
Model performance evaluation and bubble filtering. **a**. Compared with the baseline method of Yolov5, the mean average precision (mAP) of our model was high after the same number of training rounds. **b**. Comparison with original U-Net, our model produced better performance in precision, recall, and f1-score. **c**. Our model had a higher MIoU than original U-Net. d. (d1) The identification of model trained by only organoid labels. (d2) The identification of models trained by organoid labels and bubble labels. (d3). The identification of models after filtering the bubble category. **e**. A heatmap shows the accuracy of the model prediction in each training dataset. Three independent test experiments and evaluations were conducted for each identification using different methods. **f**. Average accuracy for each test evaluation in test data (n = 3 independent experiments). **g**. Guided backpropagation can indicate which pixels of organoids and bubbles in the input image contribute more to the model judgment. Grad-CAM shows important regions for predicting organoids or bubbles.

### II. Organoid Identification and Bubble Distinguish

Bubbles are inevitable in experimental operations. To exclude the effect of bubbles on organoid analysis—the first version of our network to predict and analyze organoids incorrectly identified bubbles as organoids (Fig. 3d (c1))—we conducted further analysis of the misclassification, analyzing the causes via guided backpropagation and gradient-CAM. As shown in Figure 3g, there were no obvious differences in shape and texture between organoids and bubbles in the high-level feature map, and the interest area of the network was nearly consistent. To deal with bubble misidentification, we designed two schemes: The first method uses morphological evaluation analysis to judge whether the identified organoids are air bubbles. We propose the use of bubble center and circularity evaluation to detect bubbles, because the center of a bubble is close to white, and its shape is close to a circle. The second method is to collect some bubble images and train the model with bubbles as a category, resulting in the model’s ability to identify bubbles. The identified bubbles are then filtered out and not displayed on the output picture. We randomly selected 300 images from our training data and test data several times to test whether the two proposed filtering methods effectively reduced bubbles’ misidentification as organoids. The control group was our model trained on only the organoid data. The experimental results were handed over to three experts for evaluation and calculation, and, finally, precision, accuracy, and recall were obtained (Figure 3e-f). The above two methods were determined to be useful. The work described in this paper adopted the second scheme, and its effect is shown in Figure 3d (c2, c3).

### III. Recognition of Organoid Lifetime: Growth, Fusion and Death

OrgaTracker was initialized with the result of the first organoid detection, and an ID was assigned to each organoid. However, one of the critical problems in organoid tracking is how to correlate detection objects with tracking objects. The Hungarian algorithm can be used to match and assign the IDs of organoids in adjacent frames. Since organoids grow in a hydrogel, their motion characteristics are very subtle and seldom observable; however, their growth characteristics are easy to observe and more important. Thus, ID matching and allocation were mainly based on IOU values of different organoids between frames. The organoid tracking videos were synthesized from microscope time-lapse images at different times. However, organoids varied greatly in size and area even on adjacent days (Fig. 4a). This caused the denominator to be larger when using the formula to calculate the IOU, which made the IOU value smaller. Therefore, this paper proposes a new IOU-matching criterion for organoid tracking (Fig. 4b). The denominator is the modified overlap plus the missed, which was the organoid area in the previous frame. In addition, some unmatched organoids may be newly present in the current frame, and the previous frame does not exist. According to a preset counter, OrgaTracker can assign these organoids new IDs. OrgaTracker achieved organoid tracking and area segmentation over 7 ∼ 8 d (2× magnification) (Fig. 4c). The model was also able to perform organoid tracking tasks at 4× or 10× magnification. However, tracking organoids at 2× magnification was even more challenging. More organoids were observed in the same field of view, and smaller organoids were more likely to be affected by factors such as light and lack of focus, which made their identification and area segmentation inaccurate. Therefore, we did not consider the detection and tracking of organoids that were too small. Moreover, organoid detail and texture features were less visible than those at 4× or 10× magnification. We also analyzed data for this tracking. The model was able to record organoid number, organoid area, average area, and average area change rate (Fig. 4d-f).

**Figure 4.**
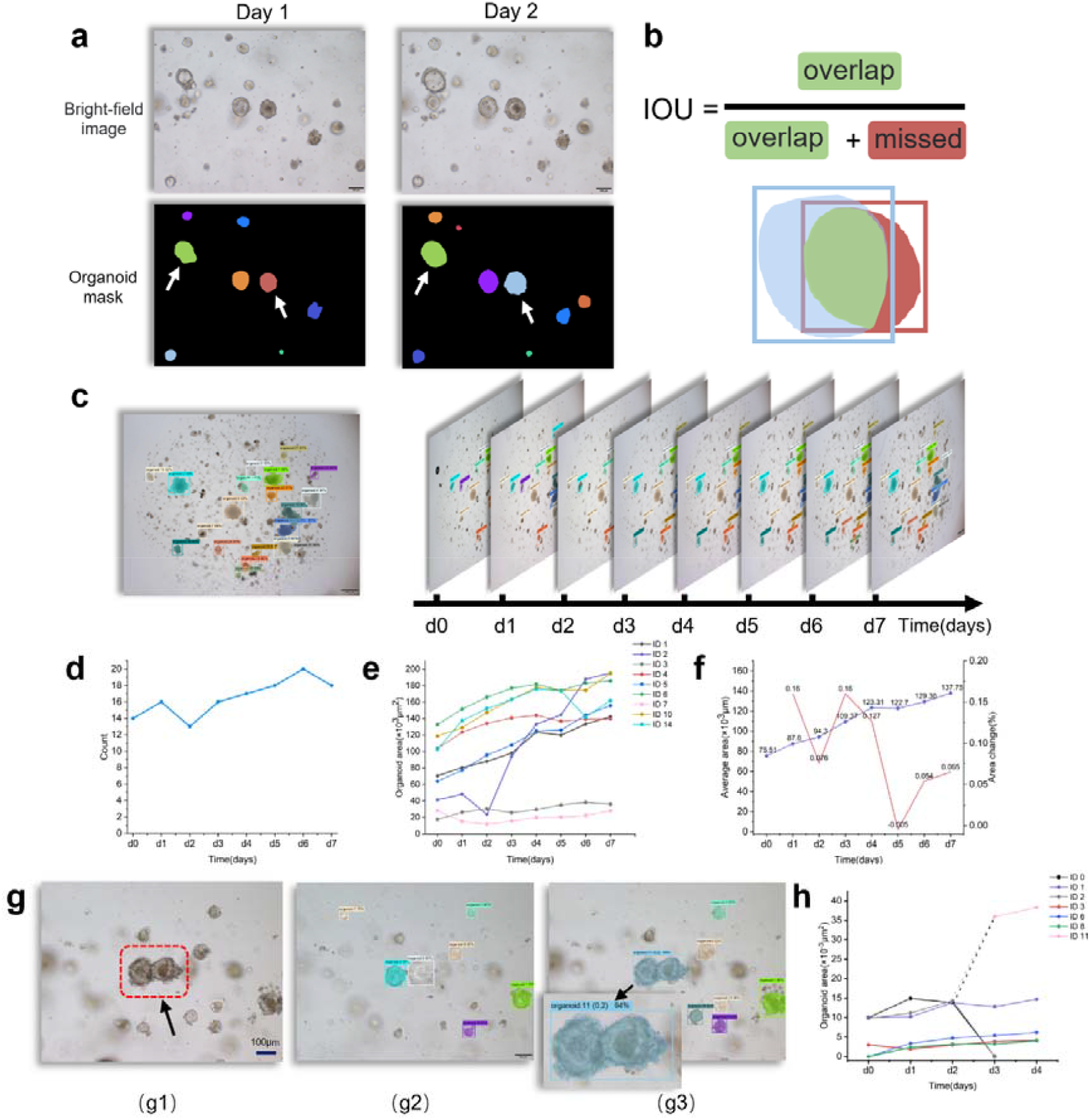
Organoid Life-time Tracking and Analysis. **a**. The size of organoids may be different at the same position in the time-lapse microscope images of different days. **b**. The improved IoU is the intersection over the smaller area of two organoid in organoid tracking. **c**. Organoids identification and segmentation over seven days. The identified and segmented organoids in time-lapse microscope images are matched across frames to realize organoid tracking over time. Shown are seven identified and segmented images sampled at 24h intervals from an organoid culture experiment. **d**. Changes in the number of organoids over seven days. **e**. The growth curves of partial organoids in Figure D were calculated over time. **f**. Organoid average area and rate of the area change in Figure E over 7 days. **g**. Organoid fusion over time. When the identified organoids are matched in adjacent frames through IoU, the algorithm calculates whether the same organoid matches multiple organoids. Moreover, the algorithm gives the fused organoid a new ID, showing which organoids it came from. **h**. The organoid area with ID 11 is formed by the fusion of the organoid with ID 0 and ID 2.

However, the traditional Hungarian algorithm can only achieve the optimal matching, i.e., one can only match one. In many cases, two or more organoids may fuse into a single organoid and form a whole (Fig. 4g (g1)). These organoids, with a low IOU, are ignored by the algorithm. Therefore, the improved algorithm traverses the matching matrix again and adds matches with an IOU greater than the threshold of 0.7 to the result set, except for the optimal matches. When an organoid in the current frame corresponds to two or more organoids in the previous frame, the model assigns a new ID to the organoid and indicates which organoids it was fused from (Fig. 4g (g2, g3)). The model accurately tracks and analyzes the area changes in organoid fusion (Fig. 4h).

We then used OrgaTracker to evaluate the effects of different clinically approved chemotherapeutics at a single dosage on human colon cancer organoids (Fig. 5a-d). Colon cancer organoids were cultured in 96-well plates for 7-8 days and injected with chemotherapeutic drugs at concentrations of 1 µm or 10 µm on day 0. The organoid image data from days 0, 4, and 7 were then taken for data analysis. The model was used to quantify the perimeter, area, roundness, and viability of the colorectal cancer organoids. Oxaliplatin, 5-fluorouracil (5-Fu), leucovorin calcium inhibited the growth of organoids, resulting in slow or reduced growth of the perimeter and area of the organoids, and reduced the viability of organoids compared with the control group. The higher the concentration of the drug, the more significant the effect. However, organoid viability was higher after treatment with 1µm 5-fu and 1µm leucovorin calcium. In addition, compared with the control group, 1µm 5-fu and 1µm leucovorin calcium reduced the roundness of organoids. High concentrations of chemotherapy drugs inhibited organoid hyperplasia, making the roundness of organoids closer to 1 compared with lower drug concentrations. However, a high concentration of oxaliplatin led to tumor organoids apoptosis, resulting in a smaller area but increased circumference and reduced roundness.

**Figure 5.**
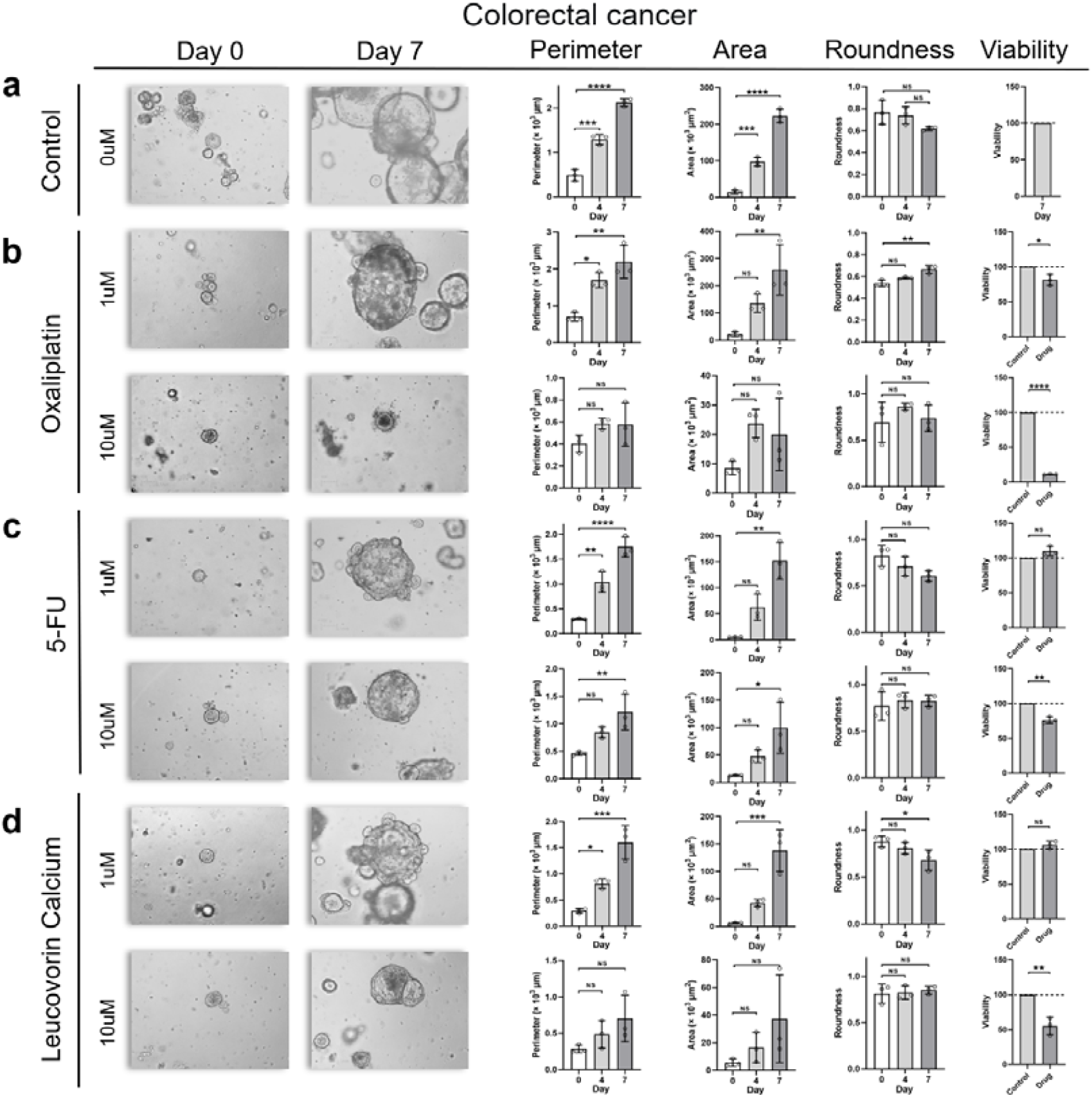
Analysis of organoid morphological changes induced by chemical drugs. **a-d**. Using deep learning algorithm to quantify the morphological characteristics of organoids after drug treatment. Organoid brightfield images (10× magnification) and quantification of the changes in perimeter, area, roundness, and viability on days 0, 4, and 7 following treatment with (a) DMSO (n = 3), (b) 1µm and 10 µm oxaliplatin (n = 3), (c) 1µm and 10 µm 5-FU (n = 3), (d) 1µm and 10 µm leucovorin calcium (n = 3).

### IV. Fine Analysis of Organoids: Budding Counting and Skeleton Extraction of Organoids

We used a two-stage object detection approach to obtain the crypt count for each intestinal organoid (Fig. 6a). First, the image was fed into the first Yolov5 network to locate each organoid. Then, the image patch of each organoid was fed into the second Yolov5 network to realize the detection and counting of small intestinal crypts. Finally, the detected information was drawn onto the original image. We trained the model using a small-intestinal organoid dataset labeled with organoids and crypts. Compared with organoids, crypts were more easily missed and misidentified by the model. Although crypts were more likely to be missed and misidentified by the model than organoids, the model still showed great performance in their identification (Fig. 6b-c). Moreover, our team conducted 50 experiments to evaluate the speed and accuracy of counting crypts between humans and the model (Fig. 6d-e). After statistical data analysis, our model was highly consistent with the human in the budding count, but much faster. We used three groups of data, namely five days of same-location organoid bright-field images, to demonstrate that the model could accurately identify and quantify crypts (Fig. 6f). In addition, the model could automatically count the number of budding organoids, the average crypt area, etc. (Fig. 6g-j).

**Figure 6.**
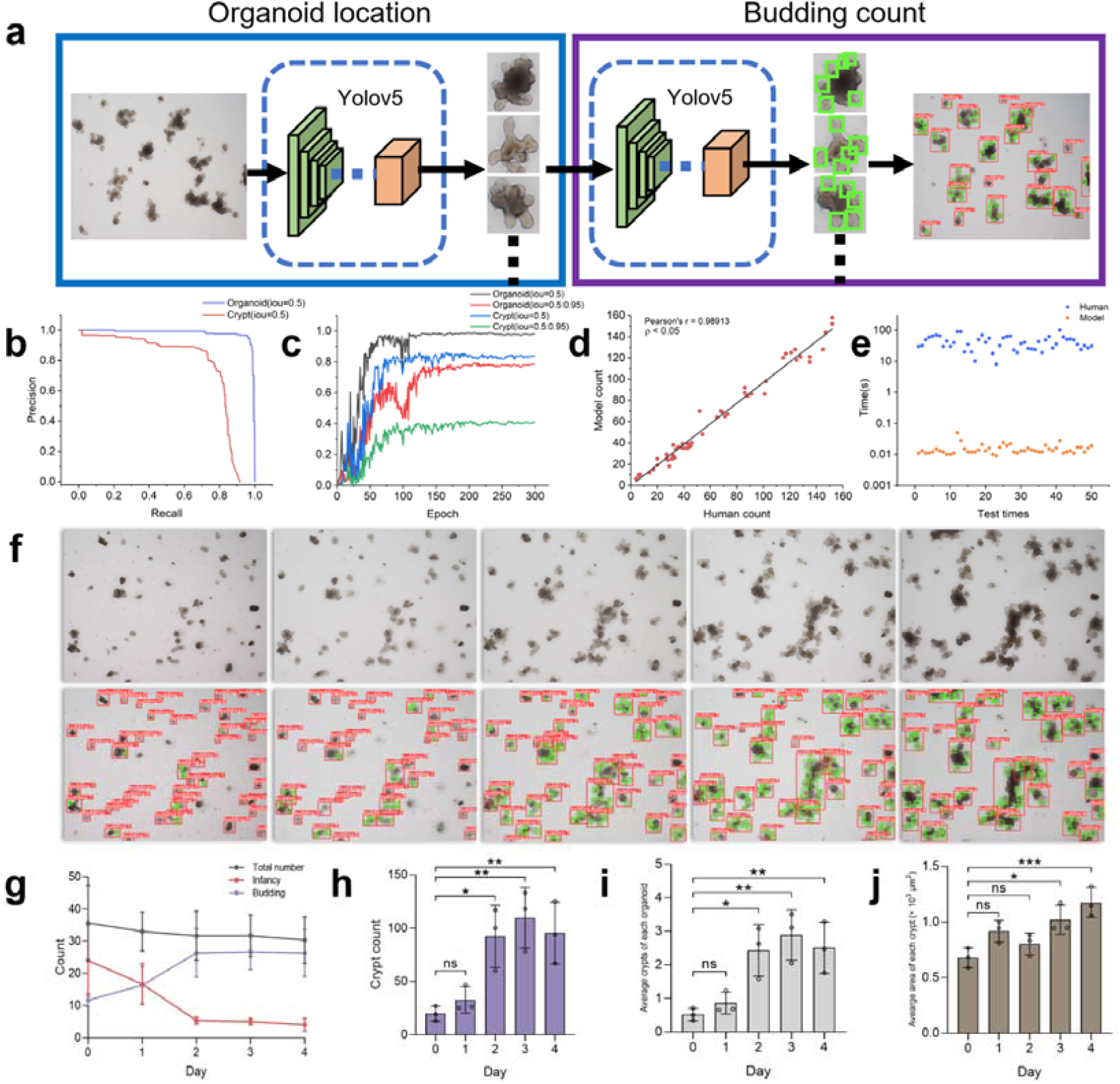
Quantification and analysis of mouse intestinal organoid budding. **a**. The network pipeline for crypt identification and counting. The input image is first sent to YOLOv5 network for organoid localization, and then the image of this region is sent to the next YOLOv5 network for the budding count. **b-c**. Evaluation of YOLOv5 network performance for organoid and crypt identification. The accuracy of organoid identification is higher than the other. Crypt identification has some errors, but the performance also meets the need. **d**. In 50 independent experiments, a comparison of the time spent by humans and algorithms to count the number of crypts in the same image. **e**. The figure shows Pearson correlations for the counting number of crypts between humans and algorithms. Through 50 independent experiments, the counting result of humans and algorithms were fitted to the line. **f**. The upper part shows five bright-field images sampled at 24h intervals from an organoid culture experiment over 4-5 d. The lower part shows the organoid localization and crypt identification of the corresponding images, and the number of budding of each organoid is shown above the organoid identification box. **g**. Shown are changes in the number of the total, budding organoids and infancy organoids over four days. **h-i**. Automatically counted the number of crypts and the average crypts number of each organoid. **j**. The area of each crypt is approximately represented by the area of the green identification box to obtain the average area of crypt.

The intestinal organoid skeleton is a helpful clue for organoid detection, complementary to the intestinal organoid contour, as it provides a structural representation to describe the relationship between absorptive villi and crypts. We used a three-stage object detection and segmentation method to achieve skeleton extraction of intestinal organoids (Fig. 7a). First, the image was fed into the Yolov5 to locate each organoid. Second, the image patch of each organoid was input into U-Net to segment the organoid and obtain its mask. Then, the organoid mask was sent to the Zhang-Suen algorithm to obtain the organoid skeleton. Last, all the information was drawn on the original image. After 100 epochs, U-Net achieved great performance in the segmentation of intestinal organoids (Fig. 7c-e). We used three groups of data, namely five days of same-location organoid bright-field images, demonstrate that the model could accurately identify and segment organoids and extract their skeletons (Fig. 7b). In addition, the model could automatically calculate the number of organoids and the average area of the organoids, extract average number of skeleton edges per organoid, and find total key points, the endpoints and intersection points in the organoid skeleton (Fig. 7f-h).

**Figure 7.**
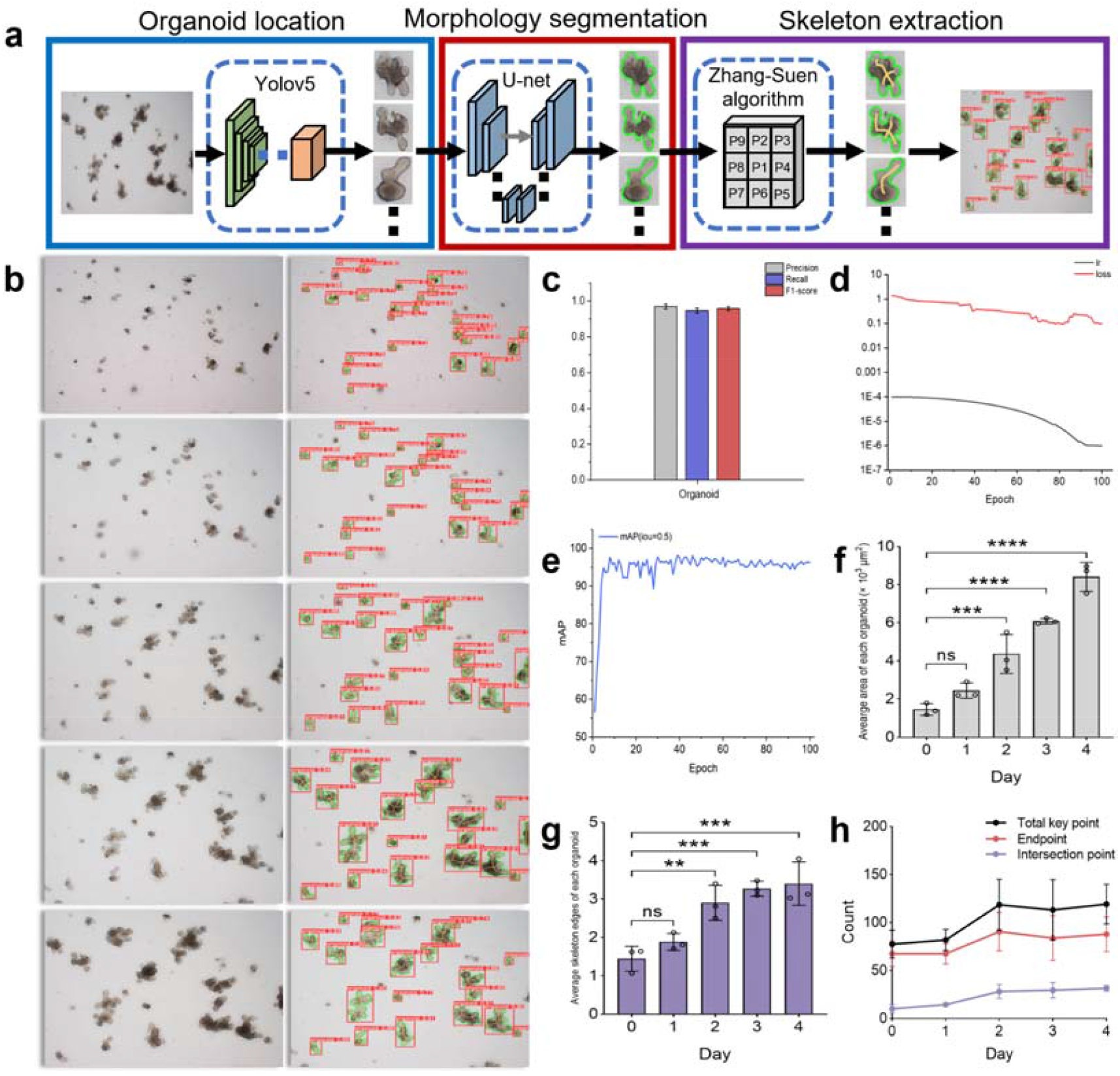
Skeleton extraction and analysis of mouse intestinal organoids. **a**. The network pipeline for skeleton extraction of mouse intestinal organoids. **b**. Shown on the left are five bright-field images sampled at 24 h intervals over a 4-5 d period in organoid culture experiments. The right part shows the organoid localization, segmentation and skeleton extraction of the corresponding images. **c**. U-Net can achieve great image segmentation of mouse intestinal organoids. **d**. After 100 rounds of training, the loss and learning rate of the network reach a small value and are in balance. **e**. After another 100 rounds of training, the mean average precision of the network is also close to a stable value. **f**. Changes in the average area of intestinal organoids in figure b over time. **g**. Changes in the average number of edges counted from the intestinal organoid skeleton over time. **h**. Changes in the number of total key points, endpoints and intersection points counted from the intestinal organoid skeleton over time.

We applied the OrgaTracker system to determine drug efficacies in the 3D intestinal organoid model using three anticancer drugs: doxorubicin (Dox), cisplatin (Cis), and paclitaxel (Pac). In addition, images are captured at the same location using z-stack during 0-4 days. We found that the total number of organoids increased or remained stable over time in the control group, while the total number of organoids remained stable or even decreased in most of the drug-treated groups. High concentrations of the drug inhibited organoid area growth compared to controls. Dox and Pac promoted an increase in the darkness of organoids, while Cis had little effect. Drugs can affect the budding of organoids, resulting in a decrease in the number of crypts in the organoids. Compared with the control group, the number of skeleton edges of organoids in the drug-treated group remained stable or gradually decreased over time, which showed that the drug prevents organoids from growing and developing, destroying their structure and making them simpler.

### IV Generation of Artificial Organoids via Sketches

Sketch-based image translation can help biologists create or design novel organoid images in practical application scenarios and is one of the most effective ways to showcase people’s creativity and exchange ideas. Image synthesis using a sketch can be considered an image translation problem in which the input is a sketch. Pix2pix is a classic work in image translation. It is based on conditional GAN; an input organoid sketch can be taken as the condition to learn the mapping between the input sketch and the output organoid image. We used and modified the pix2pix pipeline and applied it to generate various organoid images (Figure 9A). All organoids need to be sketched during model training. However, it is impractical to create sketches for all the images by hand. Therefore, the input data x was an alternative to the organoid sketch, which was the edge and internal texture of the organoid using Gaussian filtering and the Canny edge detection operator. The input data x was passed into a generator (a U-net architecture) which generated fake organoid images. The image and x formed a fake tuple, while the real original image and x formed a real tuple. The discriminator learned to classify between fake tuples and real tuples. The generator learned how to fool the discriminator and could be trained to produce more realistic organoid images after many iterations. We then trained the pix2pix pipeline using three datasets, and experts were invited to draw sketches of different organoid types. By drawing sketches of different organoids as the input, this network not only generated tumor organoids of various shapes and mouse intestinal organoids with crypt structures, but also generated complex human cerebral organoids and other images (Fig. 9b-d). The work can offer organoid images of various types for biologists to analyze and verify various organoid indicators and data. It can also provide a large amount of data for algorithms to advance the development of AI in organoids.

**Figure 8.**
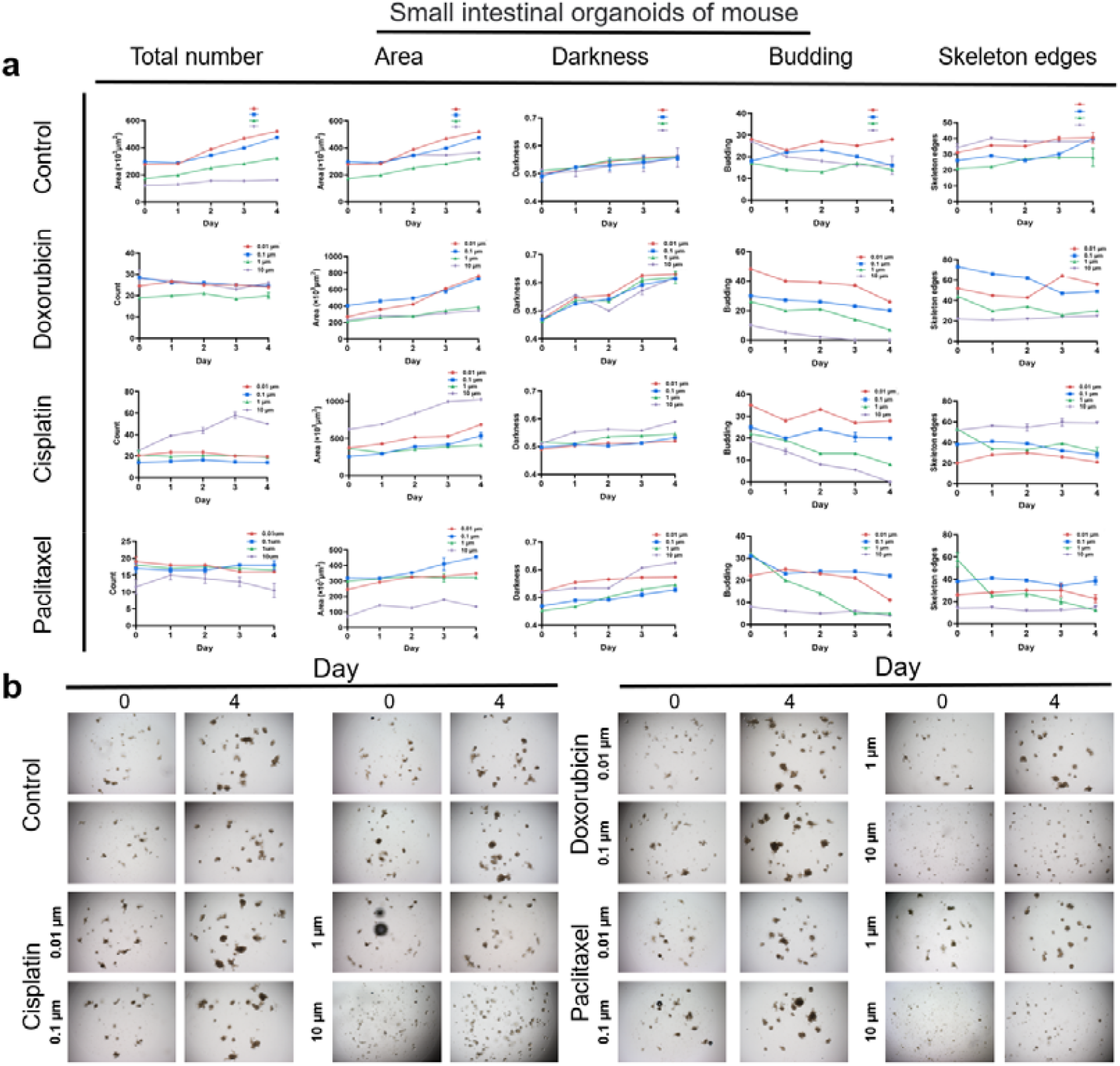
Features of intestinal organoids w or w/o drugs analyzed by OrgaTracker. a. The total number, area, darkness, budding, and skeleton edges in untreated control and drug-treated samples were analyzed and compared. b. Original data were shown. Drugs was added on day 0. Total number, area, darkness, budding, and skeleton edges were measured by the system during day 0-4. Drug concentrations used to treat each intestinal cell line were: doxorubicin, 0.01µm, 0.1 µm, 1 µm, and 10 µm; cisplatin, 0.01 µm, 0.1 µm, 1 µm, and 10 µm; paclitaxel, 0.01 µm, 0.1 µm, 1 µm, and 10 µm; Images of intestinal organoids on different days were recorded. All data are presented as mean ± S.E.M (n = 3, at least 3 repeat measures for each group).

**Figure 9.**
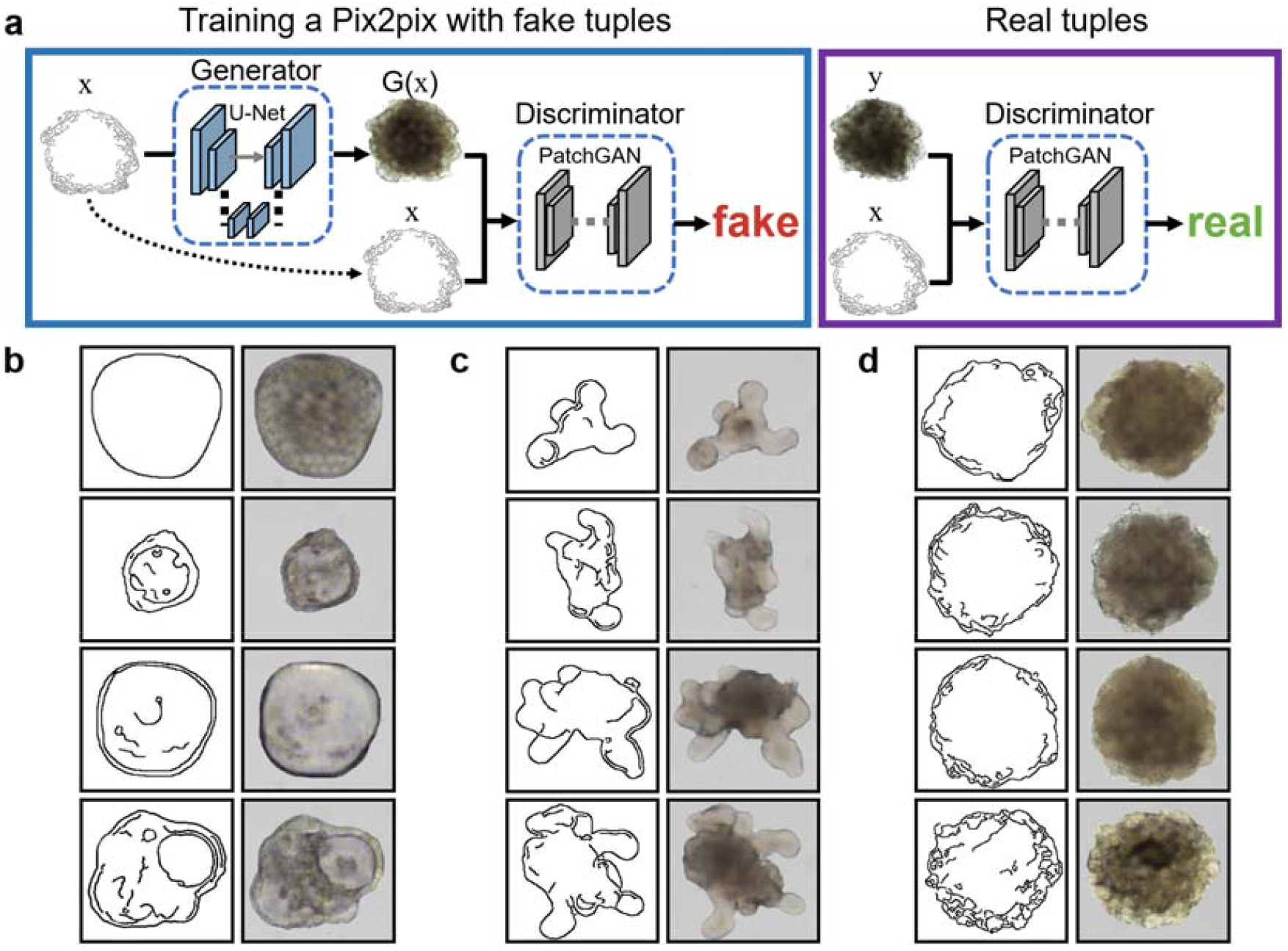
Training a Pix2pix for mapping sketch to organoid color image. **a**. Discriminator learns to classify fake tuples (formed by generator) and real (sketch, image) tuples. The generator learns how to fool the discriminator and can be trained to produce realistic organoid images. **b**. The network was used to generate tumor organoid images of different shapes using sketches. **c**. The network generated all kinds of mouse intestinal organoid images with crypt structures using sketches. **d**. The network was also capable of generating complex cerebral organoid images using sketches.

## Discussion and Conclusion

In this study, our research could be divided into three parts. Part I: We developed an organoid tracking system by combining improved target detection with a semantic segmentation algorithm. Compared with the method using only target detection, we were able to segment the organoid area in real time [11, 26]. Our models were able to track organoids at 2×, 4×, or 10× magnification. In addition, in the target detection step, bubbles were regarded as an identification class and were filtered out, which effectively avoided the misidentification of bubbles as organoids. Compared with the method that only used semantic segmentation, the method of first locating and then segmenting organoids was more accurate and also helped to avoid the mis-segmentation of overlapping or occluded organoids [27]. More importantly, the system was able to accurately capture and analyze organoid fusion (Fig. 4g and Fig. 4e) [20]. Part II: Although recent researchers have implemented deep-learning-based budding determination for human intestinal organoids, the models have only been able to distinguish between budding and nonbudding organoids, which is a classification problem. Moreover, these works used old, simple deep-learning models such as ResNet, XCeption, VGG, etc. [18]. We counted the number of the intestinal organoid crypts, and used the organoid skeleton to represent the structural relationship between intestinal villi and crypts [28]. Part III: We, for the first time, included the GAN network in organoid image generation [29, 30]. It was applied to generate images of three kinds of organoids, and outlines and detailed textures of organoids with different characteristics were generated.

However, the research still encountered some challenges. The images required for organoid tracking are microscope time-lapse images, which need to be taken from the same fixed position and collected continuously over several days [11, 31]. The process is complicated, and data collection is complex. Moreover, the growth and movement of organoids are slow, and long-term use of equipment to observe and photograph organoids may lead to a waste of resources. Therefore, it is essential to realize tracking equipment that can be automatically and intelligently cultivated and can automatically find the same location for shooting. Partially or totally out-of-focus of organoids in a 3D environment will cause great errors in long-term tracking and area calculation [10]. Using the shadows of organoids may offer an excellent method to restore out-of-focus organoid images via image restoration technology [32-34].

Cell–cell interactions affect all aspects of development, homeostasis, and disease, and cell fusion and the interactions between cells in close contact are key to health and disease [35]. Because these processes are obscure and complex, more intelligent algorithms are needed to help scientists and researchers analyze, quantify, and explain cell and organoid interactions and fusion. The number, shape, and size of mouse intestinal organoid crypts are important indicators used to evaluate whether an organoid consists of healthy growth [36, 37]. However, at present, the analysis of intestinal organoid morphology still relies on human observation, and it cannot provide further useful information. Moreover, although we have achieved budding count and skeleton analysis of mouse small-intestinal organoids in bright-field images, among other accomplishments, there is still a lack of quantitative indicators with which to evaluate intestinal organoids in other scenarios [18, 28]. Pix2pix is a universal image translation framework, in which the translation process is based on inferring the missing texture or shadow information between strokes [30, 38]. The model described in this paper used original images and the boundary edges obtained by Gaussian filtering and the Canny edge detection operator as training data [39, 40]. Only good realistic or professional sketch inputs can produce good results. This is not user-friendly for novices who lack knowledge about organoid shape and texture. Standard evaluation methods for images generated using GAN models have not been established [41]. Therefore, it may be difficult to evaluate the authenticity of the generated organoid images without the corresponding image evaluation criteria.

In conclusion, OrgaTracker can automatically track organoids and automatically segment and calculate the area of organoids while preventing interference due to bubbles, as well as capturing and analyzing organoid fusion. This model can provide indicators and analytical tools for evaluating organoids in high-throughput clinical drug screening. Our intestinal organoid analysis model provides an accurate and automatic method for quantifying small-intestinal organoid crypts and analyzing the structure of organoids, which represents great savings in manpower and material resources. Finally, we explored the application of GAN networks to organoids for the first time, and it is believed that combining these two domains will bring significant advances in organoid drug screening and other research.

## Materials and Methods

### a. Cell culture

Tumor organoid culture: we washed the tissue fragments in cold PBS containing penicillin/streptomycin (Solarbio, P1400) for 3×5 minutes, and the fragments were digested in a digestion medium containing 10 mL DMEM medium and 1% fetal bovine serum. The cells were first filtered and then centrifuged at 200 g for 5 minutes. The isolated cells were embedded in the Matrigel and placed in a preheated 24 well flat bottom cell culture plate (Costar, 3524). After the matrix ball is polymerized, add 500 μL person CRLM medium, and keep adding fresh medium every 3 days. Mouse organoid culture: we used cryopreserved mouse intestinal organoids (70931, STEMCELL Technologies) derived from the C57BL/6 mouse small intestine. According to the manufacturer’s agreement, mouse intestinal organoids were cultured in 24-well plates using IntestiCult organoid growth medium (06005, STEMCELL Technologies). In short, we add 50 µl of organoid/matrix gel mixture (1:1, v/v) into a 24-well plate to form matrix gel droplets, and then incubate it at 37°C and 5% CO_2_ for 10 min to gel the matrix gel. The preheated organoid growth medium was injected 750 μL into each well. Intestinal organoids were passaged every 4-6 d in fresh Matrigel.

### b. Drug treatment

Antitumor drugs were used at the following doses: Oxaliplatin, 5-FU, leucovorin calcium, and control (DMSO). The drug concentrations were 1µm and 10µm. The drug was dissolved according to the manufacturer’s instructions, and a 100 ×working solution was prepared with PBS buffer (Thermo Fisher).

### c. Image acquisition

The datasets contained the human colon tumor organoid dataset and the mouse intestinal organoid dataset. The first dataset consisted of 2140 organoid images (including bubble images), which was used to evaluate the proposed organoids tracking method in this paper, and the dataset was divided into two subsets with a ratio of 8:2, that is, 1926 images wASUSere used as the training and validation set, and 214 images were used as the test set. In contrast, the second dataset consisted of 600 intestinal organoids images, which were mainly used to evaluate crypt quantification and skeleton extraction of mouse intestinal organoids. Images were acquired using an Olympus IX83 motorized microscope with 2× or 4× /0.30 objective and a DFC450C camera. The growth of tumor organoids was observed continuously for 7-8 days, and the growth of small intestinal organoids was observed continuously for 4-5 days. The images were sampled every 24 hours under a bright field by manually adjusting the coordinate axes. Olympus software generated images with a resolution of 1360 × 1024. Next, each image was split into 4 images of a resolution of 400 × 400. The purpose of this operation is (1) the original image is too large, and it is easy to overflow the GPU memory. (2) the dataset can be expanded.

### d. Data annotation

Four experienced experts manually annotated all acquired images using LabelImg and LabelMe software which can easily label and save the positions and boundaries of organoids and bubbles in images. Since organoids are in 3D culture, there are likely to be out-of-focus objects under different z-axis focal planes. Out-of-focus has a great influence on automatic recognition and segmentation. However, there is no standard for judging out-of-focus except for human eyes, so we developed an auxiliary labeling tool based on the modified canny algorithm to assist experts. Each image was annotated by two experts, and if the agreement was below 80% (as determined by calculations of Intersection over Union (IoU)), the image was submitted to a third expert for further annotation.

### e. Data preprocessing

Each image was normalized according to its standard deviation. In addition, to improve the generalization and anti-interference ability of the model, the image geometry transformation and image intensity modification make the data expand. This method can be used to prevent model overfitting and allow models to perform better with smaller initial datasets.

### f. Video generation

The video consists of microscope time-lapse images of organoids cultured in the same well plate for several days. Using Python library functions, multiple images were synthesized as MP4 video at one frame per second.

### g. Algorithms for organoid tracking and analysis

#### g1. Improved Yolov5 architecture

Yolov5 consists of input, backbone network, neck network, and detection head [42, 43]. The input side represents the input image. Yolov5 uses Mosaic [17] data enhancement operation to improve the training speed of the model and the accuracy of the network and proposes adaptive image scaling processing to scale the image to a uniform size. The backbone Network is made up of a CNN and a transformer. The neck network consists of Feature Pyramid Networks (FPN) and Path Aggregation Networks (PAN). The output of the detection head still adopts CIoUs as the loss function of the Bounding box and provides several detection scales. Finally, DIoU-NMS is used to suppress the target detection box by non-maximum value to obtain the optimal prediction box.

#### g2. Bottleneck transformer block

Traditional CNNs have translational invariance and efficient capture of local features, but lack the ability to model globally and over long distances. The transformer can make up for them. The bottleneck transformer block replaced the 3×3 convolutional layer of the ResNet bottleneck block with Multi-Head Self-Attention [42, 43].

#### g3. The Improved non-maximum suppression

Distance Intersection over union-non-maximum suppression (DIoU-nms) considers not only the IoU, but also the distance between the center points of two boxes [44]. When the IoU between the two boxes is large, but the distance between the two boxes is also far, it may be considered that they are two objects and will not filter out one of them.

#### g4. Nested U-Net architecture

U-Net is an asymmetric network composed of multiple encoders and decoders. It consists of convolution, down-sampling, up-sampling, and concatenation operations. The compression path of this network consists of four blocks, each containing two 3×3 convolutions, two RELUs, and a maximum pooling. The compression path can extract important image features and reduce image resolution. The extended path of this network also includes four blocks, each with two 3×3 convolutions, two RELUs, and one up-sampling. In addition, the feature map of each layer needs to be merged with the feature map of the symmetric compression path on the left. The extended path can combine deep and shallow features to get better segmentation results. We added a series of nested skip pathways between encoders and decoders, which aim to narrow the semantic gap between the feature maps of the encoder and decoder sub-networks [24, 45].

#### g5. Organoid area calculation

We use the scale in the image to calculate the length represented by each pixel to get the area represented by each pixel and count the number of pixels in the mask output by U-Net. Multiplying them together will get the real area of the organoid.

#### g6. The improved intersection over union

The intersection of the two bounding boxes in two adjacent images ratio the area of the bounding box in the previous image.

#### g7. Mask optimization

Acquired organoid masks output by U-Net were processed analysis of the connected domain, and the largest connected domain was preserved in each mask. The largest connected domain was used to avoid incorrect segmentation of organoids.

#### g8. Analysis of small intestine organoid budding

The budding number refers to the number of crypts per organoid, and the budding rate is the average number of crypts per organoid in the same field of view. The area of the green bounding box approximately represents the area of the crypt.

### h. Bubble classification of artificial characteristics

#### Bubble center pixel intensity

We first calculate the center point of the bounding box, and then get the average value of the 2*k*+1◊2*k*+1 region with it as the center point.

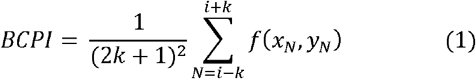

Where (*x*_*i*_, *y*_*i*_), *f*(*x*_*N*_,*y*_*N*_,), and *k* are the center point of the bounding box, RGB value of the point, and half the length of the region without considering the center point, respectively.

#### Bubble circularity

Bc is defined as the ratio between the object area and the object perimeter. For a perfect circle, this ratio will approach the value of 1.

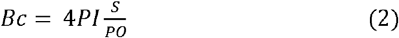

Where S and PO are the area, and the perimeter, respectively.

### i. Guided backpropagation and grad-CAM

Guided backpropagation was used to show which pixels of bubbles and organoids are of interest to a convolutional neural network, and Grad-cam was used to visualize important regions of bubbles and organoids [46, 47].

### j. Evaluation of network performance

Network performance was evaluated by recall, precision, accuracy, F1 score and mean average precision, and the AUC of the ROC curve. Recall refers to the proportion of positive examples that are predicted to be correct in the entire sample. Precision represents the proportion of true positive samples among predicted positive samples. Accuracy is the proportion of samples that are predicted to be correct out of the total sample. F1-score is the harmonic average of precision and recall. *MAP* is the average AP value, which is calculated from multiple validation sets. *MIoU* is the average of intersection over union of each class in the dataset, where *i* represents the true value, j represents the predicted value, and *P*_*ij*_ represents the predicted value of i as j.

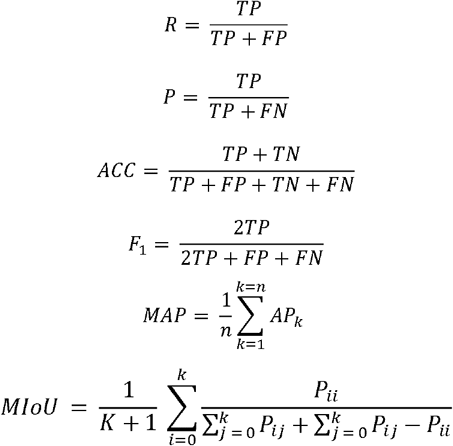

### k. GPU server and analysis environment

We used a GPU server, which has two CPUs: Xeon 4-Core E5-2637V4 3.5 GHz, 128 GB CPU memory, and four GPUs, GeForce GTX1080Ti GDDR5 11GB (NVIDIA, Santa Clara, CA, USA). We programmed all scripts on the Nvidia-docker system with Ubuntu 18.04, CUDA 1.10.1, cuDNN 11.3, Anaconda3 4.4.0, Python 3.8.2, Pytorch 1.10.0.

## Data and materials availability

All data required to evaluate the conclusions of the paper have been provided in the paper and/or Supplementary Materials. Additional data related to this paper may be requested from the authors. The custom code can be available in the public GitHub repository.

## Acknowledgement

This work was supported by the National Key R&D Program of China (2017YFA0700500) and National Natural Science Foundation of China (Grant No. 62172202), Experiment Project of China Manned Space Program HYZHXM01019, and the Fundamental Research Funds for the Central Universities from Southeast University:3207032101C3.

## Author contributions

XD, ZZC, and YHL were involved in conceptualization; XD, JPS, and WHC contributed to formal analysis; JJG, LFS, and ZZG were involved in funding acquisition; XD and WHC helped in investigating; YPC, MLL, QWL, and JZ helped in drug testing. XD, JPS, YXQ, and WHC contributed to image annotation; YHL and ZZG were involved in project administration. ZZC and YHL helped in supervision; XD, JPS, and WHC contributed to validation; XD, ZZC, and YHL were involved in writing original draft; XD, ZZC, and YHL helped in writing-review & editing.

## Declaration

### Ethical approval

This study does not contain any studies with human or animal subjects performed by any of the authors.

